# Discontinuation of GLP-1RA Treatment Reinstates Binge-Like Eating and Drives Body Weight Past Pre-Treatment Levels in Mice

**DOI:** 10.64898/2026.11.02.763781

**Authors:** Britny A. Hildebrandt, Susanne E. Ahmari

## Abstract

We examined the impact of semaglutide treatment and discontinuation in a pre-clinical model for binge eating. Treatment reduced body weight and palatable food intake in mice with binge-like eating. Discontinuation reinstated binge-like eating and caused body weight to significantly exceed pre-treatment baseline, revealing specific binge-related risk when discontinuing GLP-1RAs.

## Introduction

Use of glucagon-like peptide-1 receptor agonist medications (GLP-1RAs, e.g., semaglutide) has risen dramatically in recent years [1], with evidence highlighting the successful use of GLP-1RAs to treat a range of medical and psychiatric conditions (e.g., type II diabetes, obesity, alcohol use disorder [2,3]). Growing evidence also suggests that GLP-1RAs may reduce binge eating (BE) [3,4], a widely prevalent dysregulated eating behavior characterized by the intermittent overconsumption of highly palatable food (PF) [5]. Despite this promise, GLP-1RAs are not currently FDA-approved for BE, and the mechanisms through which they impact BE remain poorly understood [6]. Furthermore, discontinuation of GLP-1RAs is common in clinical practice. A large proportion of individuals discontinue use within a year, often due to cost or unwanted side effects [7], and discontinuation is associated with significant weight regain (e.g., 60-90%) within one year [8]. This highlights an urgent need to understand the physiological and behavioral consequences of GLP-1RA use and discontinuation on eating patterns and weight, particularly in vulnerable BE populations. However, little is known about the behavioral patterns of dysregulated eating after GLP-1RA discontinuation, particularly in individuals with a history of BE. The rapid increase in GLP-1RA medication use and discontinuation, combined with unknown effects on individuals with BE, has raised a critical and clinically relevant issue: does relapse following discontinuation of GLP-1RAs differ in individuals with and without a history of BE? Identification of rebound BE following GLP-1RA use could inform clinical management of GLP-1RA treatment and discontinuation in groups at risk for eating disorders.

As a first step towards identifying neural mechanisms underlying GLP-1RA response, we used an established animal model of BE [9] to map the full behavioral trajectory of food intake and weight changes during GLP-1RA treatment and discontinuation. This mouse model allows comparison of a BE group, in which intermittent, binge-like access to PF produces escalating PF intake in otherwise normal-weight animals, and a continuous access binge-control group (CO), in which continuous PF access produces weight gain without a binge-like intake pattern. Using this framework, we tested the hypothesis that semaglutide treatment would suppress PF intake and weight in BE and CO mice, but that discontinuation would selectively reinstate binge-like eating in BE mice. This pattern of results would identify a clinically urgent relapse risk specific to individuals with a history of BE and establish this model for future investigation of underlying circuitry.

## Methods

Eighteen adult female C57BL/6J mice (Jackson Laboratories) were randomized to either intermittent binge-like eating (BE) or continuous control (CO) access to palatable food (PF) and completed the paradigm for BE [9] before undergoing 14 days of semaglutide treatment and a 14 day discontinuation phase. Detailed methods are provided in Supplemental Materials. All experiments were approved by the Institutional Animal Use and Care Committee at the University of Pittsburgh in compliance with National Institutes of Health guidelines for the care and use of laboratory animals.

## Results

Two-way repeated measures ANOVA using phase-averaged PF intake (i.e., average PF intake across the final three days of each phase: pre-treatment, semaglutide treatment, discontinuation) showed a significant Group x Time interaction (Fig. 1A, F(2, 32) = 13.74, p<.0001). PF intake decreased from the pre-treatment to the treatment phase in the BE group (p<.0001). PF intake in BE mice rebounded to pre-treatment levels during discontinuation with no significant difference between pre-treatment and end of the discontinuation phase (p=.9992), showing recurrence of pre-treatment binge-like eating behavior. In contrast, CO mice showed no difference in PF intake across phases (all ps>.10). Day-by-day analysis of PF intake confirmed phase-averaged patterns such that PF intake differed by group (F(14, 224) = 2.258, p=.0068), with BE mice suppressing PF intake on all 14 semaglutide treatment days (Fig. 1B, all ps<.0001). During discontinuation, day-by-day analysis also showed a significant Group x Time interaction (Fig. 1C, F(14, 224) = 3.485, p<.0001). BE mice rebounded to pre-treatment levels of PF intake, showing no significant differences from pre-treatment (Fig. 1C, ps>.10) apart from three timepoints (days 2, 4, 7, all ps =.0047-.0292). CO showed no differences from pre-treatment baseline PF intake at any day across treatment and discontinuation phases (all ps >.10).

**Fig. 1.**
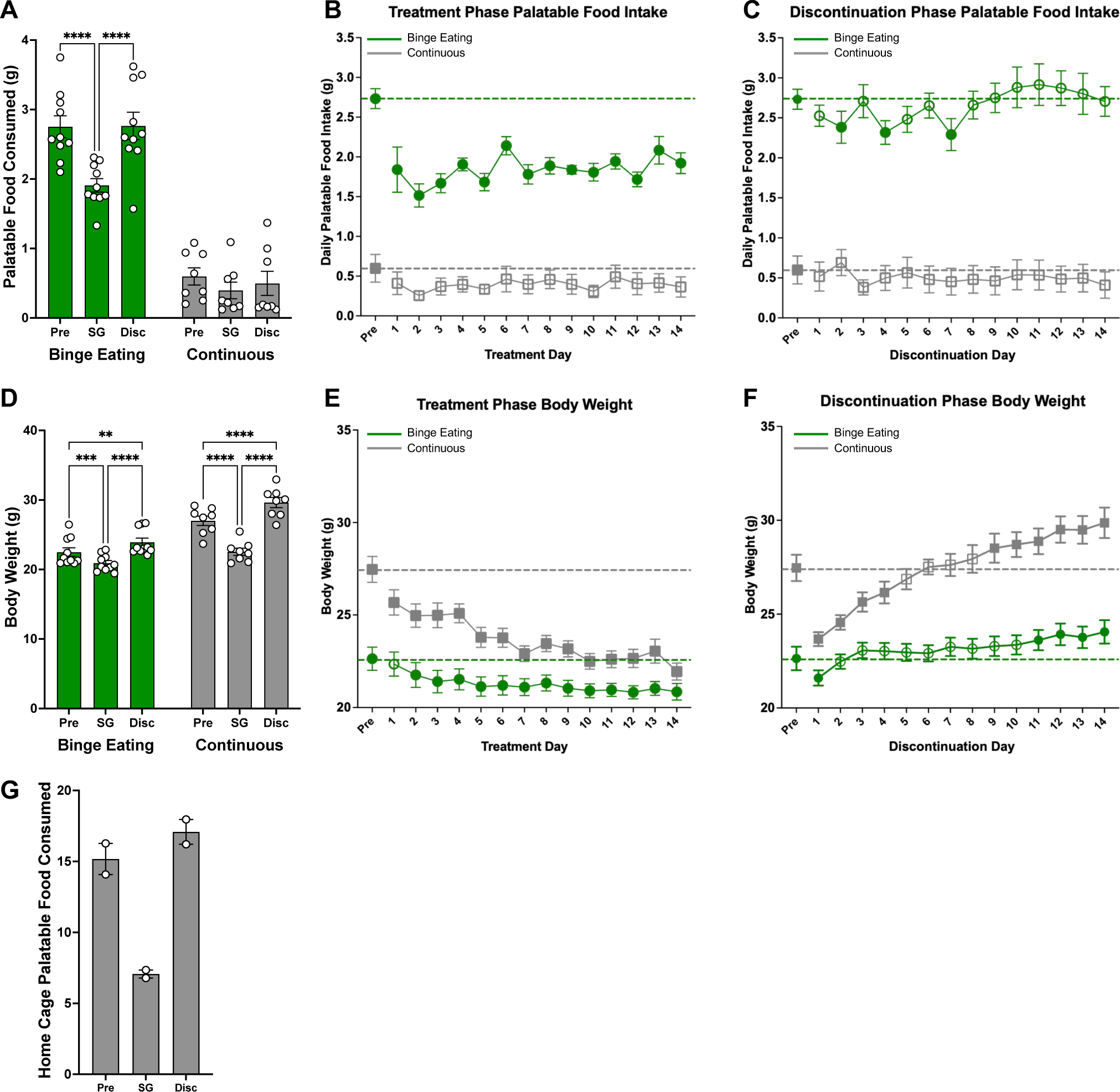
Effects of semaglutide treatment and discontinuation on palatable food intake and body weight. A) 2-hour Palatable food (PF) intake averaged across the last 3 days of each phase (pre-treatment [Pre], semaglutide treatment [SG], discontinuation [Disc]) in Binge Eating and Continuous mice. Daily 2-hour PF intake across the B) treatment and C) discontinuation phases. D) Body weight (BW) averaged across the last 3 days of each phase (Pre, SG, Disc). Daily BW across the E) treatment and F) discontinuation phases. Dashed lines in B, C, E, F indicate each group’s pre-treatment baseline. G) Continuous mice 22-hour home cage PF intake averaged across the last 3 days of each phase (n = 2 home cages, 4 mice/cage) (Pre, SG, Disc). No statistical test performed due to small home cage sample size; provided for illustrative value. Data are mean ± SEM. BE n=10, CO n=8. In A and D, data were analyzed by two-way repeated-measures ANOVA with Šídák’s post hoc tests. In B, C, E, F, data were analyzed by two-way repeated-measures ANOVA with Dunnett’s post hoc comparisons to pre-treatment PF or BW. Filled symbols indicate significant difference from pre-treatment baseline (Dunnett’s, p<.05); open symbols indicate non-significant comparisons. *p<.05, **p<.01, ***p<.001, ****p<.0001.

Analysis of phase-averaged weight also showed a significant Time x Group interaction (Fig. 1D, F(2,32)=31.25, p<0.0001). Both groups had significantly decreased weight in the treatment phase (BE p=.0002, CO p<.0001). Critically, both BE and CO mice regained weight during the discontinuation phase and significantly exceeded their pre-treatment baseline (BE p=0.0010; CO p<0.0001). Day-by-day analyses using two-way repeated measures ANOVA showed that weight differed by group during treatment (Fig. 1E, F(14,224)=9.266, p<0.0001) and discontinuation (Fig 1F, F(14,224)=21.28, p<0.0001). Post-hoc tests indicated CO lost significant weight every treatment day, and BE did so from treatment day 2 onward. During discontinuation, both groups returned to pre-treatment baseline weight before significantly exceeding pre-treatment weight. Specifically, CO mice exceeded pre-treatment weight starting discontinuation day 9, and BE mice starting discontinuation day 11.

## Discussion

GLP-1RA use and discontinuation has significantly accelerated in recent years. While there is some evidence for GLP-1RA effectiveness for treating BE behavior, there is currently a significant gap in our understanding of the mechanisms that contribute to the effects, hindering development of appropriate clinical interventions to manage both GLP-1 treatment and potential BE relapse during discontinuation. Pre-clinical approaches are uniquely positioned to model pharmacological treatments while observing impacts on behavior and neural circuitry. This study aimed to establish a model of the full treatment and discontinuation trajectory of semaglutide using an animal model for BE to pave the way for future studies probing underlying mechanisms.

Findings showed that BE mice rebounded to pre-treatment binge-like levels of PF intake within one day of GLP-1RA discontinuation. Furthermore, weight in both BE and CO groups rebounds quickly following GLP-1RA discontinuation. Critically, weight continued to increase during discontinuation, exceeding pre-treatment weight in both groups. This pattern of effects raises an important concern for individuals with a history of BE specifically–i.e., quick return to dysregulated eating patterns, combined with the physiological weight gain beyond pre-treatment levels seen during discontinuation and unknown changes in the neural circuitry governing eating behavior, raises concern for the clinical impacts of GLP-1RA use in individuals with BE. While BE PF intake rapidly returned to binge-like levels without exceeding pre-treatment baseline within this two-week window, it is possible that intake could exceed baseline (as observed for weight) if a longer discontinuation period was used. Together, these data highlight an important challenge: the eating disorders field has not yet reckoned with the reality that GLP-1RAs are increasingly prescribed to and discontinued by individuals with BE, and clinicians currently have limited evidence to anticipate whether, when, or how eating behavior and weight may evolve during this vulnerable period. Furthermore, this problem likely extends well beyond those with diagnosed BE, as BE is also common among individuals seeking obesity treatment [10] but often goes undiagnosed [11]. Thus, many patients currently receiving GLP-1RAs through obesity care may be at unrecognized risk.

The observed dissociation between PF intake returning to pre-treatment levels, while weight exceeded pre-treatment, suggests that feeding behavior and energy balance may recover along different trajectories following GLP-1RA discontinuation. Weight continued to increase over baseline after PF intake had returned to pre-treatment levels, likely reflecting additional contributions from energy expenditure or metabolic rate beyond intake alone. This suggests that feeding behavior alone may be an incomplete proxy for physiological recovery following discontinuation. In CO mice, the two-hour PF measure showed no significant change from baseline at any point in treatment or discontinuation, despite robust weight effects, consistent with reduced sensitivity of this limited-access measure in a phenotype with continuous home-cage PF availability. Supporting this interpretation, CO home cage (22-hour) PF intake data declined from pre-treatment phase average of 15.18 grams/cage to 7.07 grams/cage during the treatment phase before rebounding to 17.08 grams/cage during discontinuation (Fig 1G), a pattern that paralleled the excessive weight gain beyond pre-treatment observed in this group. Because all CO mice were housed across only two cages (n = 4/cage), these home cage data reflect cage-level consumption rather than individual intake, precluding formal statistical comparison. These descriptive data nonetheless provide evidence suggesting that the behavioral response of CO mice to GLP-1RA discontinuation may be considerably larger than the two-hour measure alone would indicate. Future work using this model should incorporate more precise measurements of individual animal home cage intake to better analyze feeding trajectories in the CO group.

This paradigm establishes a model for identifying the neural circuits underlying differential recovery following GLP-1RA discontinuation in BE. However, findings are limited to a single semaglutide concentration/dose, a 14-day discontinuation window, and a single sex (female). Whether findings would generalize to different GLP-1RA doses/compounds, longer treatment/discontinuation phases, or male mice is unknown. Nevertheless, because reward and homeostatic feeding circuitry implicated in BE substantially overlap with circuitry underlying other dysregulated, reinforcement-based behaviors (e.g., alcohol and substance use), which have also shown positive impact from GLP-1RA medications [2], our findings may extend beyond eating disorders to have broader transdiagnostic implications. Together, our results suggest that discontinuation of GLP-1RAs could affect relapse vulnerability in disorders characterized by reward dysregulation, highlighting an urgent priority for future research.

