## Supplemental Materials for "Discontinuation of GLP-1RA Treatment Reinstates Binge-Like Eating and Drives Body Weight Past Pre-Treatment Levels in Mice"

Britny A. Hildebrandt, Ph.D.^1,2^ & Susanne E. Ahmari, M.D., Ph.D. ^1,2,3^

^1^Department of Psychiatry, University of Pittsburgh School of Medicine, Pittsburgh, PA, USA, 15213

^2^Translational Neuroscience Program, Department of Psychiatry, University of Pittsburgh, PA, USA, 15213

^3^Center for Neuroscience, University of Pittsburgh, Pittsburgh, PA, USA, 15213

*Animals*. Eighteen adult female C57Bl/6J mice (Jackson Laboratories, 8-10 weeks of age at start of study) were used. Mice were group housed with 2-4 mice per cage and had *ad libitum* access to food and water throughout the study. Animals were maintained on a 12/12-hour light-dark cycle (lights off at 7:00 AM; on at 7:00PM). All experiments were approved by the Institutional Animal Use and Care Committee at the University of Pittsburgh in compliance with National Institutes of Health guidelines for the care and use of laboratory animals. Experimenters were not blinded to group assignment (BE, CO) during the study, as feeding condition was necessarily apparent based on each animal’s home cage and feeding test cage setup.

*Binge Eating Paradigm*. The current study used a well-established paradigm for BE [1,2] which uses different intermittent access schedules to palatable food (PF) to elicit binge-like eating in rodents. All mice were randomized to one of two feeding groups, ensuring for no baseline weight differences: 1) binge eating (BE), in which animals received binge-like intermittent access to PF daily for 2-hours; or 2) continuous control (CO), in which animals received continuous (24-hour), non-intermittent access to PF (sweetened condensed milk [Nestle; 3.25 kcal/g; 22% fat, 67% sugar, 10% protein; diluted in a 3:1 ratio with water]). All mice had *ad libitum* access to standard chow (Prolab Isopro 5P76; 4.1 kcal/g; 14% fat, 60% carbohydrates, 26% protein) throughout the paradigm and were never placed on food restriction.

Mice completed five weeks of daily PF feeding tests starting at approximately 2-3 hours after dark (active phase) onset. Prior to each test, all mice were removed from group-housed conditions and placed in individual test cages for the duration of each 2-hour test. PF (delivered via 50mL conical tube with sipper attachment) and standard chow were weighed at the beginning and end of the 2-hour test, and body weight was measured daily. At completion of each test, mice were removed from test cages and returned to group housed home cages. The PF tube in the CO animals home cage was weighed daily before the feeding test and replaced. All paradigm feeding test procedures continued across the treatment and discontinuation phases for a total of 49 feeding tests (35 initial BE paradigm, 14 treatment days, 14 discontinuation days).

*Semaglutide treatment.* Following completion of the paradigm for BE, all mice received daily subcutaneous injections of semaglutide (0.1 mg/kg; semaglutide acetate, MedChemExpress [Cat. Number: HY-114118B], Monmouth Junction, New Jersey) dissolved in sterile saline. Semaglutide dosing was calculated daily based on each animal’s measured body weight from the same day to account for any weight changes over the course of treatment. Injections were administered daily, approximately 3-4 hours following completion of the daily feeding test to minimize potential confounding effects of treatment-related malaise on feeding behavior. Mice received semaglutide for 14 consecutive days (treatment phase), followed by a 14-day discontinuation phase during which no further drug was administered.

*Statistical Analyses.* All statistical analyses were performed in GraphPad Prism (version 10.6.1). Sample sizes were n=10 Binge Eating (BE) and n=8 Continuous access (CO) mice (N=18 total); no missing data occurred throughout the full study. Phase-averaged PF intake and body weight were calculated as the mean of each animal’s final three days within each phase (Pre-treatment, Semaglutide, Discontinuation). Two-way repeated-measures ANOVA, with Phase (Pre-treatment, Semaglutide, Discontinuation) and Group (BE, CO) as factors, were used to examine phase-averaged PF intake and weight. Significant interactions were followed by Šídák’s post-hoc tests to evaluate within-group differences across phases and between-group differences within each phase.

Day-by-day PF intake and weight were analyzed using two-way repeated-measures ANOVA, with Study Day and Group as factors, run separately for the treatment phase (14 days) and discontinuation phase (14 days). Within each phase, planned comparisons of each day against the pre-treatment baseline day (i.e., the day immediately preceding the initiation of semaglutide treatment) were conducted using Dunnett’s multiple comparisons test, run separately within each group (BE, CO).

Home cage (22-hour) PF intake data for CO mice were collected at the cage level (n=2 cages, 4 mice/cage) and are therefore presented descriptively (mean values per phase) rather than analyzed inferentially, given the insufficient number of independent units for statistical comparison.

For all analyses, α was set at 0.05, and sphericity was assumed for repeated-measures ANOVAs as implemented in Prism. Exact p-values are reported throughout except where p < 0.0001.
